# 17 β-estradiol impedes aortic root dilation and rupture in male Marfan mice

**DOI:** 10.1101/2023.05.09.540071

**Authors:** Louis Saddic, Sean Escopete, Lior Zilberberg, Shannon Kalsow, Divya Gupta, Mansoureh Egbhali, Sarah Parker

**Author notes:** Corresponding author: Sarah J Parker PhD, Assistant Professor, Cardiology, Smidt Heart Institute, Co-Director, Proteomics and Metabolomics Core Cedars Sinai Medical Center, Room 223.6 North Campus Innovation Center 8687 North Melrose Avenue Los Angeles, CA 90069. Funding for this work includes: 1R01HL165471-01(Parker), R00HL128787, Bethesda MD (Parker), and the FAER mentored research training grant, Schaumburg, IL(Saddic).

## Abstract

Marfan syndrome causes a hereditary form of thoracic aortic aneurysms with dilation of the aortic root. Human and animal models suggest a worse phenotype for males compared to females with respect to aneurysm size and risk of dissection. In this study we examine the effects of 17 β-estradiol on aortic dilation and rupture in a Marfan mouse model. Marfan male mice were administered 17 β-estradiol and the growth in aortic root size along with the risk of aortic rupture or dissection with the addition of angiotensin II was measured. Transcriptomic profiling was used to identify enriched pathways from 17 β-estradiol treatment. Aortic smooth muscle cells were then treated with cytokines in order to validate the mechanism of 17 β-estradiol protection. We show that 17 β-estradiol decreased the size and rate of aortic root dilation and improved survival from rupture and dissection after treatment with angiotensin II. The Marfan transcriptome was enriched in inflammatory genes and the addition of 17 β-estradiol modulated a set of genes that function through TNFα mediated NF-κB signaling. These included many proteins known to play a role in the phenotypic shift of aortic smooth muscle cells from a contractile to a more inflammatory-like state such as Vcam-1, Mcp-1, Lgals3, Il-6, Il-1b, and C3. In addition, 17 β-estradiol suppressed the induction of these TNFα induced genes in aortic smooth muscle cells in vitro and this effect appears to be NF-κB dependent. In conclusion, 17 β-estradiol protects against the dilation and rupture of aortic roots in Marfan male mice through the inhibition of TNFα -NF-κB signaling and thus prevents the phenotypic switch of aortic smooth muscle cells from a contractile to an inflammatory state.

## Introduction

Thoracic aortic aneurysms (TAAs) are among the most dangerous types of cardiovascular disease due to the high morbidity and mortality associated with acute dissection.^1^ Epidemiological studies have consistently demonstrated sex disparities among patients with TAAs with a higher prevalence in men (1.7:1).^2, 3^ Similar disparities exist for abdominal aortic aneurysms (AAA). Animal models of AAA have shown that this protection may be driven by 17 β-estradiol as mice prone to AAA and treated with 17 β-estradiol have smaller abdominal aortic diameters while similar mice undergoing ovariectomy have the opposite effects.^4, 5^ The etiology of AAA is considered mechanistically distinct from that of TAA which is highlighted by the fact that many forms of TAA are due to heritable gene mutations. Unfortunately, very little is known regarding mechanisms underlying sex bias in aneurysm and dissection severity in heritable forms of TAA. Among Marfan patients, despite equal probability of inheriting FBN1 mutations, males are more likely than females to have aortic root dilation and dissection.^3, 8, 9^ Likewise, studies in the Marfan *Fbn1*^*C1039G/+*^ mouse have shown that male *Fbn1*^*C1039G/+*^ mice have larger aortic root diameters compared to females after correction for body size.^6, 7^ ^3, 8, 9^ In this same model, inhibition of androgen receptors by flutamide treatment in male mice attenuates TAA growth in part through blocking transforming growth factor beta (TGFβ) signaling.^10^ Interestingly, it has been shown that flutamide treatment can enhance circulating 17 β-estradiol levels which was not examined in prior work.^11, 12^ Despite a clear role for 17 β-estradiol in protection from AAA, parallel studies to examine the potential therapeutic effects of 17 β-estradiol in *Fbn1*^*C1039G/+*^ mice or any other model of heritable TAAs have yet to be performed. A clear understanding of mechanisms conferring protection due to female sex could contribute to therapeutic interventions for both males and females that will allay disease severity and risk for dissection.

The protective effects of 17 β-estradiol against vascular injury have been studied in many contexts including hormonal regulation of extracellular remodeling and atherosclerosis.^13^ Many of these therapeutic actions have been linked to 17 β-estradiol’s ability to curtail inflammation. Specifically, 17 β-estradiol has been shown to attenuate both the attraction of inflammatory cells and the upregulation of inflammatory markers following vascular injury.^14^ Increased inflammation has been documented in degenerative forms of AAAs and TAAs in the presence of atherosclerosis.^15^ Nevertheless, inflammation may also play a role in many forms of hereditary and syndromic forms of TAAs affecting younger patients including those in Marfan syndrome. Aortic tissue from Marfan patients have increased infiltration of macrophages, B cells, and CD4+ and CD8+ T cells.^16^ In addition, fibrillin-1 fragments from mice may have the ability to attract macrophages directly.^17^

In this study we demonstrate the effects of 17 β-estradiol on the development and rupture of aortic aneurysms in the Marfan mouse model *Fbn1*^*C1039G/+*^. We provide evidence that 17 β-estradiol downregulates TNFα stimulated NF-κB pro-inflammatory genes in vivo and in smooth muscle cells in vitro. These findings provide insight into potential mechanisms for the observed sex differences in the prevalence and outcomes of patients with TAAs including Marfan syndrome and could unmask novel therapeutic targets for both sexes.

## Methods

### Mice

All animal procedures were approved by the Cedars-Sinai Medical Center Institution Animal Care and Use protocol. Male and female wild-type or *Fbn1*^*C1039G/+*^ Marfan mice were maintained on a 129 genetic background with >20 back-crossings. 17 β-estradiol treatment was accomplished through anesthetizing 8-week old mice with continuous isoflurane gas. The dorsum of mice were shaved and sterilized with betadine prior to making a sub-centimeter subcutaneous incision in order to place a 17 β-estradiol pellet (0.25mg/pellet 60 day release, Innovative research of America, Sarasota, FL). Sham surgery was performed on littermate controls. Mice were imaged for aortic root and ascending aorta size with transthoracic echocardiography under anesthesia every 2 weeks from the date of surgery until animal sacrifice with isoflurane gas 8 weeks later. For aortic rupture studies, 8-week old mice were treated with 17 β-estradiol pellet or control surgery for 4 weeks followed by implantation of subcutaneous mini-osmotic pumps (Azlet, Cupertino, CA) for the continuous infusion of angiotensin II (1000ng/kg/min). Surgical implantation of pumps was conducted in a similar matter described for pellet placement. Mice were aged for an additional 1 week for tissue harvesting and RNA expression analysis or 4 weeks for survival analysis.

### Histology

After animal sacrifice, mice were perfused with 10cc of cold phosphate buffer saline through the left ventricle. The heart was transected across approximately the atrioventricular groove and the ascending aorta, arch, and distal thoracic aorta was dissected from the mouse. The heart and aortic tissue were then placed in 4% paraformaldehyde for 24 hours at 4°C and then transitioned to 70% ethanol. Fixed tissue was embedded in paraffin and sectioned at the aortic root for hematoxylin and eosin, trichome, and elastin staining. Images were taken using the ECHO Revolve microscope (ECHO, San Diego CA).

### Western Blot

After animal sacrifice, aortic root tissue was dissected from the aortic valve to the sinotubular junction and then immediately flash frozen in liquid nitrogen. Protein lysates from RIPA buffer were reduced in loading buffer, run on acrylamide gels and transferred to polyvinylidene fluoride membranes. The membranes were then probed with antibodies against Mmp2 and Mmp9 (Cell Signaling, Danvers MA) and Vinculin (Abcam, Waltham MA). Quantification was performed on ImageQuant™ LAS4000 (GE) with the use of Vinculin as a loading control.

### Cell Culture

Aortic smooth muscle cells were extracted from four different male Marfan mice. Cells were never passaged beyond eight times. Cells were grown in phenol red free DMEM/F-12 with 10% charcoal stripped serum until 90% confluent. Cells were then serum starved and pre-treated with 17 β-estradiol (1 mM, Sigma, Waltham MA) or vehicle for 24 hours followed by 8 hours of recombinant TNFα (2ng/ml, R&D Systems, Minneapolis MN). Similar experiments were conducted with pre-treatment of cells for 2 hours with Pyrrolidine Dithiocarbamate (PDTC, 25mM, Sigma, Waltham MA) or vehicle followed by 8 hours of TNFα (2ng/ml) or vehicle.

#### RNA Isolation and Quantitative Real-Time PCR

Cells or aortic root tissue were harvested in TRIzol (Thermo Fischer, Waltham MA) and RNA was purified with phenol/chloroform. Reverse transcriptase (Applied Biosystems, Waltham MA) followed by quantitative real time PCR (qPCR) was then performed with SYBR green (Bio-rad, Hercules CA). Ct values for RNA transcripts were normalized to *Gapdh* expression using the delta delta Ct method. Statistical significance was determined using a Kruskal-Wallis test. The primer sequences used include mouse: Gapdh f:GGCATTGCTCTCAATGACAA r:ATGTAGGCCATGAGGTCCAA, C3 f:ACCTTACCTCGGCAAGTTTCT r:TTGTAGAGCTGCTGGTCAGG, Mcp-1 f:GCATCCACGTGTTGGCTCA r:CTCCAGCCTACTCATTGGGATCA, Vcam-1 f:TGAACCCAAACAGAGGCAGAGT r:GGTATCCCCATCACTTGAGCAGG, Il-6 f:TGAACAACGATGATGCACTTG r:CTGAAGGACTCTGGCTTTGTC.

### RNA-seq

RNA from mouse aortic roots was extracted as described above. Libraries were prepped and sequencing was performed on the Illumina HiSeq3000 (Illumina, San Diego, CA). Sequencing depth was roughly around 30 million reads per sample. Raw reads were analyzed for quality using FASTqc^18^. Reads were aligned using TopHat2 version 2.1.1^19^ and reads were counted using HTseq version 2.0.2^20^. Differential expression and principle component analysis was performed with DESeq2^21^ and pathway analysis was conducted with the gene set enrichment algorithm (GSEA)^22^. Transcription factor target enrichment was performed on the TRUUST database from the Enrichr bioinformatic platform^23^. Plots were made in R studio using ggplots2^24^. Significant differentially expressed genes used the cutoff adjusted p-value <0.1 and absolute fold change >1.5. Adjusted p-value for multiple comparison uses the Benjamini Hochberg method per default settings of DESeq2.

### Statistics

Aortic root size, ascending aorta size, mouse weight measurements, and in vitro RNA expression data are expressed as mean +/-standard error. Descriptive statistics for aortic size quotients includes median and interquartile range (IQR). Kruskal-Wallis is used to describe statistical significance between groups. Survival significance was determined using the Wilcoxon method. Graphs and statistical analysis were performed in JMP (Cary, NC) or R studio^25^.

## Results

### *17* β*-estradiol attenuates aortic root enlargement in Marfan mice*

The aortic root was measured by transthoracic echocardiography in 8-week old mice every two weeks for a total of 8 weeks. Male Marfan mice treated with 17 β-estradiol for 4, 6, and 8 weeks had significantly smaller aortic root diameters compared to littermate male Marfan mice treated with placebo. In fact, the mean aortic diameter of 17 β-estradiol treated males tended to trend closely to the mean aortic diameter of littermate female Marfan mice, which itself was also significantly smaller than that of male Marfan mice. Female Marfan mice treated with 17 β-estradiol also tended to have decreased root diameters comparted to littermate female Marfan mice but this difference was not statistically significant (Figure 1A). Aortic root growth, as calculated by the quotient of aortic root diameter at 16 weeks of age over the aortic root diameter at baseline, was significantly higher in the Marfan male mice (median 1.22 IQR 1.1-1.4) compared to Marfan male mice treated with 17 β-estradiol (median 1.11 IQR 1.1-1.1) (p-value = 0.00047). There was a trend towards higher aortic root growth in female Marfan mice (median 1.15 IQR 1.1-1.2) compared to those treated with 17 β-estradiol (median 1.09 IQR 1.1-1.2) but this was not statistically significant (p-value = 0.27) (Figure 1B). There was no significant differences in the change in ascending aorta diameter or weight over the duration of the 8-week study (Supplemental Figures 1A and 1B). Despite changes in root size on gross inspection (Figure 1C), Marfan male mice and littermates treated with 17 β-estradiol both had intact intimal layers and similar degrees of elastin breaks (Figure 1D). Treatment with17 β-estradiol did, however, result in reduced levels of two well established molecular markers of aneurysm pathology, matrix metalloproteinase-2 (Mmp2) and matrix metalloproteinase-9 (Mmp9), in male Marfan mice (Figure 1E).

**Figure 1.**
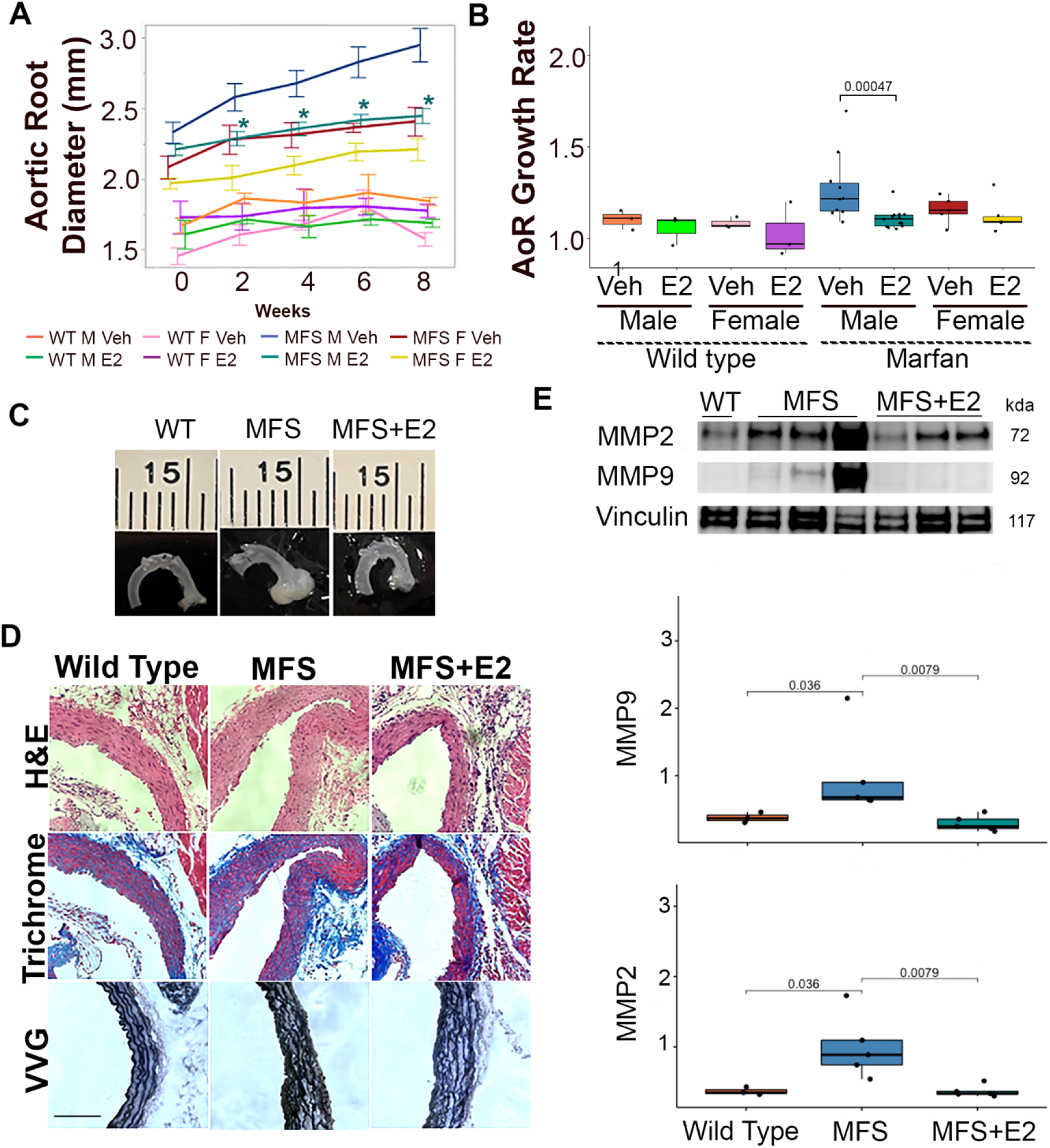
17 β-estradiol attenuates aneurysm growth in male Marfan mice. A.) Aortic roots measured every 2 week from wild-type male (n=3), wild-type female (n=3), wild-type male + 17 β-estradiol (n=3), wild-type female + 17 β-estradiol (n=3), Marfan male (n=10), Marfan male + 17 β-estradiol (n=13), Marfan female (n=5), and Marfan female + 17 β-estradiol (n=6) mice. Data are mean +/-standard error. * denotes p<0.05. B.) Ratio of aortic root diameter at 8 weeks over baseline. Data are median and interquartile range. Mouse cohorts as in A. C.) Representative aortic images from male wild-type, Marfan, and Marfan + 17 β-estradiol mice. D.) Representative H+E, trichome, and elastin sections. Mouse cohorts as in C. Scale bar=90mm. E.) Western blot with quantification of aortic root lysates from male wild-type (n=3), Marfan (n=5), and Marfan + 17 β-estradiol (n=5) mice probed for matrix metalloproteinases. E2: 17 β-estradiol, MFS: Marfan syndrome, Veh: vehicle, WT: Wild-type.

### 17 β-estradiol prolongs survival in an aortic rupture mouse model

Despite significant dilation, histological disorganization, and progressive growth in Marfan aortic roots, unchallenged *Fbn1*^*C1039G/+*^ Marfan mice rarely dissect or rupture. Therefore, we employed the established approach of an angiotensin II challenge to provoke a more consistent phenotype severity, promote progression to dissection/rupture, and evaluate the potential beneficial effect of 17 β-estradiol^26^. We observed 100% death within 30 days in male Marfan mice treated with angiotensin II. Female Marfan mice treated with angiotensin II had a significantly higher rate of survival compared to males (p=0.02) as did Marfan male mice treated with 17 β-estradiol (p=0.01). Angiotensin II had no effect on the survival of wild-type male or female mice, who exhibited 100% survival after 30 days of Angiotensin II infusion (Figure 2A). On gross examination, the majority of Marfan mice died from aortic rupture in various locations including the aortic root, the ascending aorta, the aortic arch, the descending thoracic aorta, and the abdominal aorta (Figure 2B). Those Marfan mice without evidence of rupture had signs of aortic root dissection on histological examination. Marfan mice treated with 17 β-estradiol had an equal percentage of dissected aortas but less rupture and more mice without evidence of rupture or dissection (Figure 3B).

**Figure 2.**
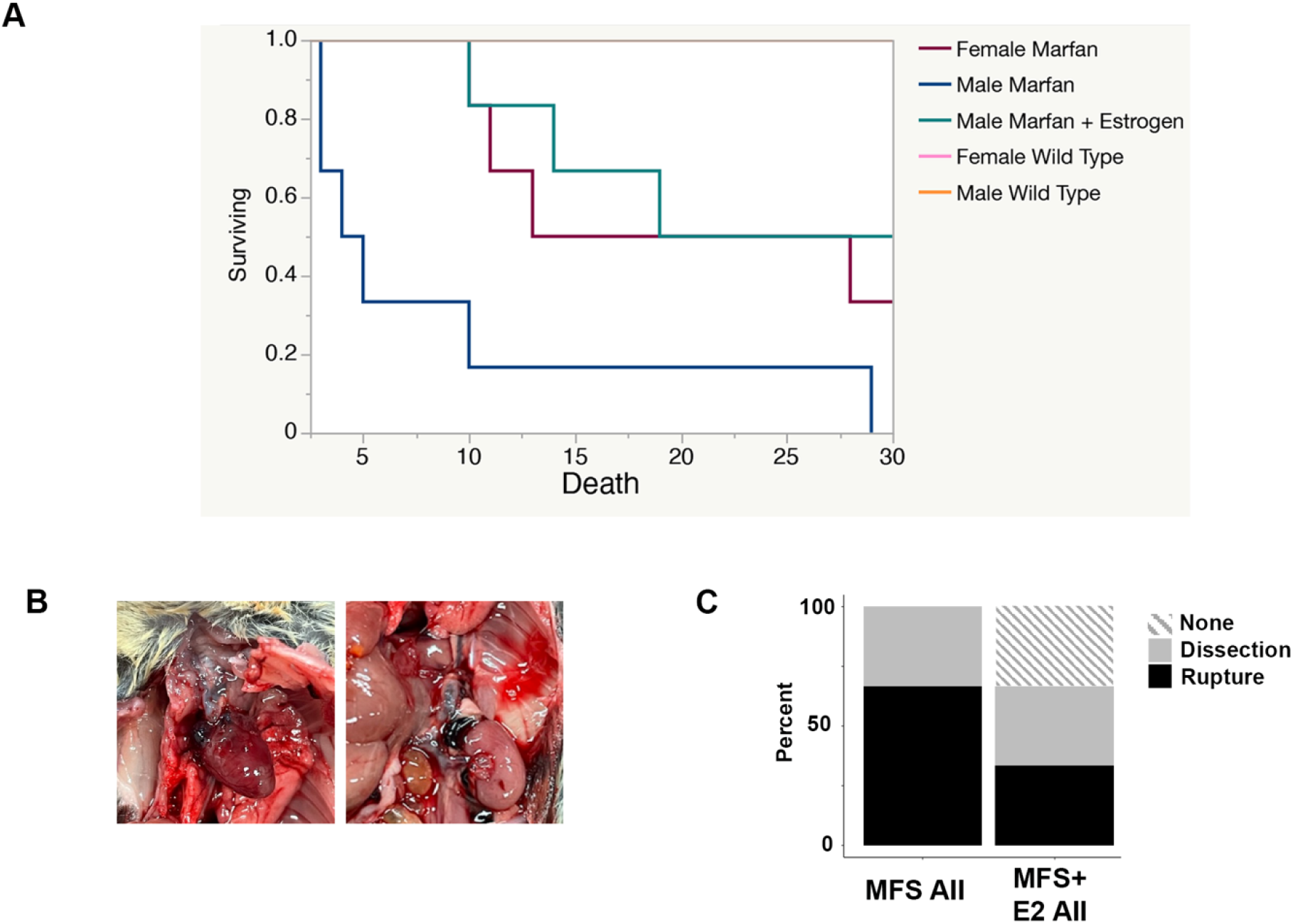
17 β-estradiol protects against aortic rupture in Marfan male mice treated with angiotensin II. A.) Survival curve of Marfan male, Marfan female, wild-type male, wild-type female, and Marfan male mice pretreated with 4 weeks of 17 β-estradiol following continuous infusion of angiotensin II. B.) Representative pictures of thoracic (left) and abdominal (right) aortic rupture in Marfan male mice treated with continuous infusion of angiotensin II. C.) Percentage of aortic pathologies found on gross and histological examination of aortas from angiotensin II treated Marfan males in the presence or absence of 17 β-estradiol. AII: angiotensin II, E2: 17 β-estradiol, MFS: Marfan syndrome.

**Figure 3.**
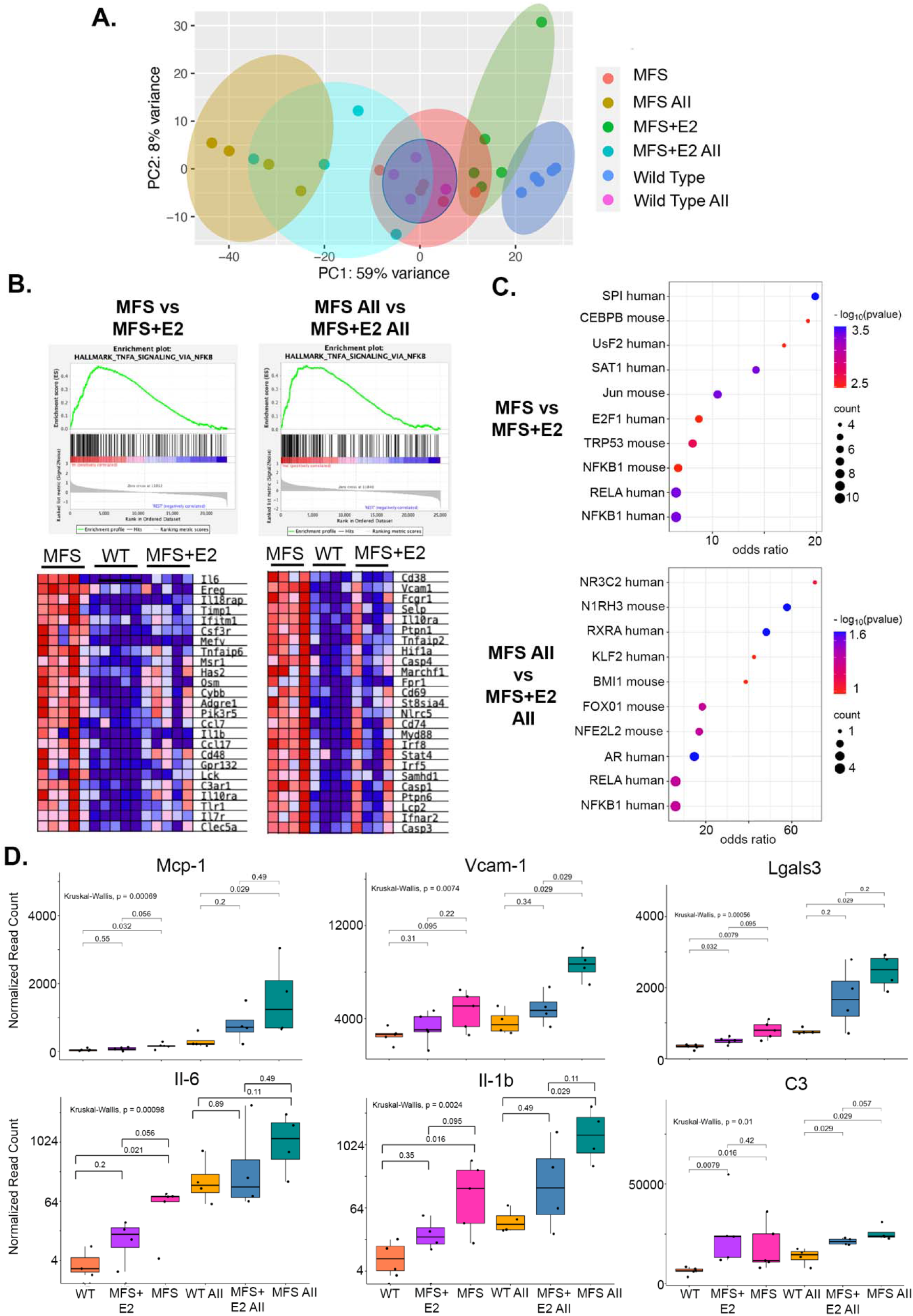
Transcriptomic profiling indicates that 17 β-estradiol blocks TNFα -NF-κB signaling and inflammation A.) Principle component analysis plot from RNAseq performed on aortic root tissue of male wild-type (n=5), Marfan (n=5), Marfan treated with 8 weeks of 17 β-estradiol (n=5), and 7 days of angiotensin II treated wild-type (n=4), Marfan (n=4), and Marfan mice treated with 5 weeks of 17 β-estradiol (n=4). B.) Gene set enrichment analysis of the hallmark gene set TNFα signaling via NF-κB comparing Marfan mice to a cohort of combined wild-type and Marfan mice treated with 17 β-estradiol (left). The same analysis in angiotensin II treated mice (right). C.) Dot plot of transcription factors with targets enriched in Marfan mice compared to Marfan mice with 17 β-estradiol using the TRRUST database (top). The same analysis in angiotensin II treated mice (bottom). D.) Normalized read counts of *Mcp-1, Vcam-1, Lgasl3, Il-6, Il-1b*, and *C3* gene expression from the same mouse cohorts outlined in A. Box plots describe quartiles. p-values for pairwise and group comparisons are depicted above charts. AII: angiotensin II, E2: 17 β-estradiol, MFS: Marfan syndrome, WT: wild-type.

### 17 β-estradiol Attenuates an Inflammatory Molecular Phenotype in Male Marfan Mice

To gain insight into potential underlying mechanisms of 17 β-estradiol protection we performed transcriptomic profiling on aortic roots from littermate 16-week old male wild-type (n=5), Marfan (n=5), and Marfan mice treated with 8 weeks of 17 β-estradiol (n=5). We also profiled aortic roots from 13-week old male wild-type, Marfan, and 17 β-estradiol treated Marfan mice exposed to continuous infusion of angiotensin II for 7 days, a timepoint just before death begins significantly increasing in Angiotensin II treated male Marfan mice. Principle component analysis demonstrated that global gene expression changes of Marfan aortic roots and wild-type aortic roots treated with angiotensin II diverge from wild-type tissue by a similar magnitude.

This divergence is even more extreme in the Marfan male mice treated with angiotensin II. 17 β-estradiol pulls the global gene expression closer to the wild-type in both the angiotensin II treated and non-angiotensin II treated cohorts (Figure 3A). Using gene set enrichment analysis (GSEA) we compared Marfan mice to a combined cohort of wild-type and 17 β-estradiol treated mice in the presence and absence of angiotensin II and observed significant enrichment in the hallmark gene set TNFα signaling via NF-κB (Figure 3B). Similarly, differentially expressed genes upregulated in Marfan mice and suppressed with 17 β-estradiol in the presence and absence of angiotensin II were enriched in NF-κB target genes in the TRRUST database of human and mouse transcription factors (Figure 3C). Among the most differentially expressed of these NF-κB target genes are proteins known to play a role in phenotype switching of aortic smooth muscle cells (SMCs) to a more inflammatory-like state and include monocyte chemoattractant protein-1 (*Mcp-1)*, vascular cell adhesion molecule-1 (*Vcam-1)*, Galectin 3 (*Lgals3)*, interleukin-6 (*Il-6)*, interleukin-1beta (*Il-1b)*, and complement C3 (*C3)* (Figure 3D).

Similarly, an opposite trend existed with SMC contractile markers such as *aSMA*, transgelin (*Tagln*), myosin heavy chain 11 (*Myh11*), and calponinin (*Cnn1*) which were downregulated in Marfan mice but to a lesser degree in those Marfan mice treated with 17 β-estradiol (Supplemental Figure 2).

### 17 β-estradiol inhibits TNFα mediated NF-κB target gene expression in Marfan aortic smooth muscle cells

Previous studies have shown that 17 β-estradiol can block TNFα induced NF-κB target genes in in many different cell lines including rat aortic SMCs^27^. In order to determine if the gene expression changes in NF-κB target genes uncovered in our bulk RNA-seq experiments could be taking place in SMCs and establish that 17 β-estradiol can directly modulate these effects we performed in vitro experiments on aortic SMCs harvested from four independent male Marfan mice. Stimulation of these cells with TNFα induced the expression of the NF-κB target genes *Mcp-1, Vcam-1, Il-6*, and *C3*. Pre-treatment with 17 β-estradiol significantly inhibited the induction of these genes by TNFα (Figure 4A). In order to demonstrate that 17 β-estradiol mediated these effects through NF-κB, these same aortic SMCs from Marfan mice were treated with TNFα in the presence or absence of the NF-κB inhibitor PDTC. As expected, PDTC blocked the TNFα mediated induction of *Mcp-1, Vcam-1, Il-6*, and *C3* (Figure 4C).

**Figure 4.**
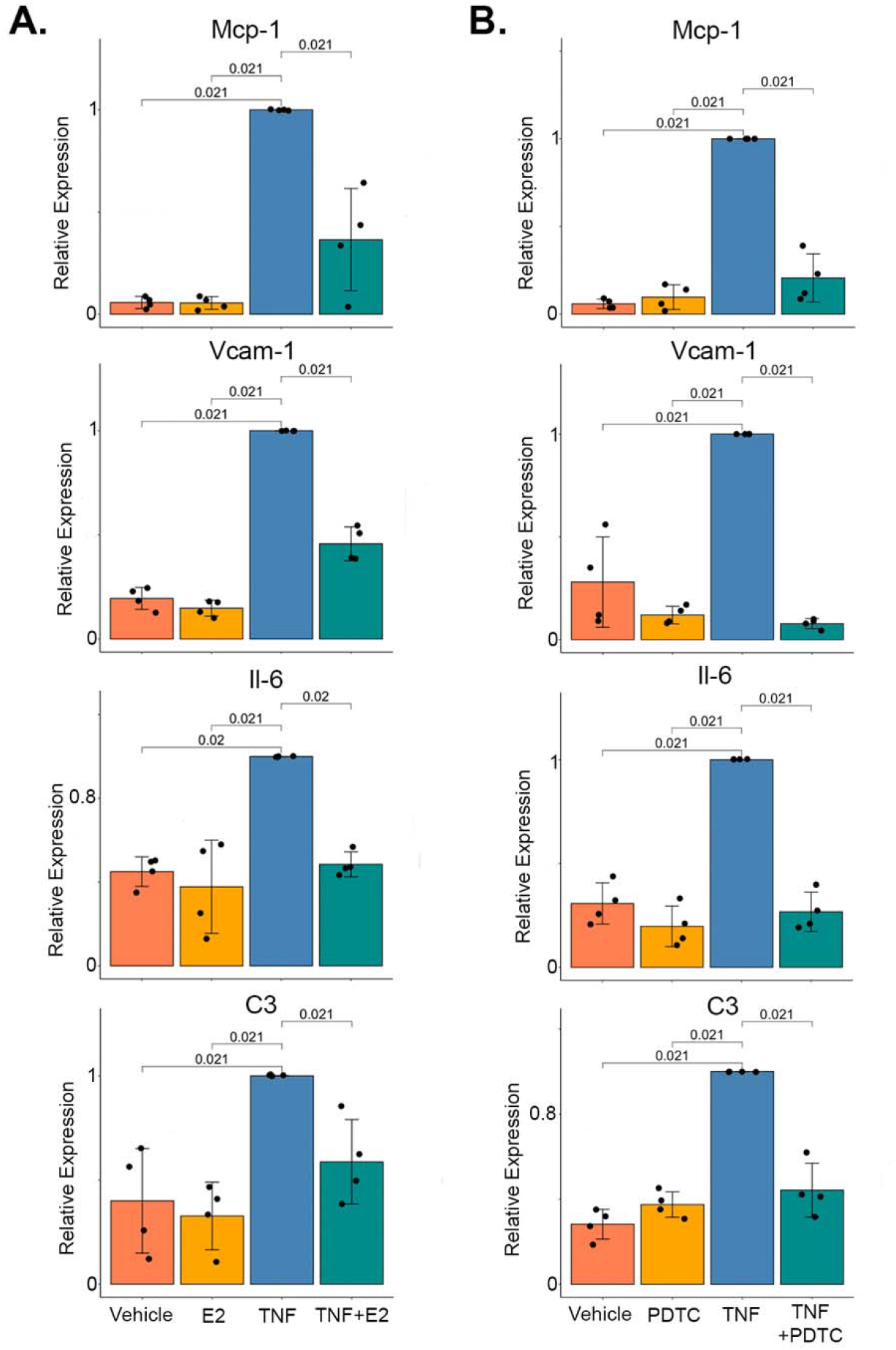
17 β-estradiol mitigates TNFα medicated NF-κB gene expression in aortic SMCs. A.) Expression of *Mcp-1, Vcam-1, Il-6, and C3* in n=4 independent Marfan male aortic SMC lines pre-treated with 17 β-estradiol or vehicle followed by the addition of TNFα or vehicle. B.) Expression of genes as in A. following pre-treatment with PDTC or vehicle followed by the addition of TNFα or vehicle. Bar charts display mean +/-standard error. Relative expression is normalized to the cytokine or drug alone treated group. E2: 17 β-estradiol, MFS: Marfan syndrome, PDTC: Pyrrolidine Dithiocarbamate, SMC: smooth muscle cell.

## Discussion

In this study we show for the first time that 17 β-estradiol treatment attenuates aortic root growth and reduces angiotensin II mediated aortic rupture in a mouse model of hereditary TAA. These findings are consistent with prior evidence supporting 17 β-estradiol mediated protection in AAA^5, 28^, and extends this knowledge to the etiologically and mechanistically distinct yet to date unexamined hereditary TAA. Interestingly, female mice treated with additional 17 β-estradiol also trended toward having a slower progression of aortic root growth compared to female mice treated with vehicle, but this effect was substantially less prominent compared to males, not statistically significant, and did not fully return female Marfan root dimensions to wild-type levels. The reduction in aneurysm size correlated with reduced expression of matrix metalloproteinases Mmp2 and Mmp9. 17 β-estradiol has been shown to both directly regulate expression of these extracellular matrix proteins and interact with several upstream molecular pathways that converge on the control of their expression.^29^ Our data are consistent with prior investigations of male-biased TAA severity in Marfan syndrome, however previous work has focused on androgen signaling^10^. Notably, these studies have shown that androgen receptor inhibition by flutamide attenuated aortic root growth, which at face value suggests that our results with 17 β-estradiol may complement or act via similar, directionally opposed mechanisms relative to androgens. Our observation that, while potently reducing aortic dimensions, 17 β-estradiol did not reverse or completely stop growth of aortic roots through the 8-week time course, is consistent with this possibility. Importantly though, as mentioned above, androgen receptor inhibition has been shown to elevate circulating 17 β-estradiol levels,^11, 12^ which was not examined in the previous study. This observation, coupled with our data indicate that protection from aneurysm growth subsequent to flutamide treatment may be at least somewhat mediated by a direct 17 β-estradiol-related effect. Certainly there remains significant room for additional studies to better understand how sex hormones impact bias in aneurysm and dissection severity in Marfan and other hereditary aortopathies.

Translational utility of the knowledge that 17 β-estradiol protects against aneurysm progression in males will rely upon a more defined molecular mechanism downstream of global hormone therapy, as estrogen supplementation to treat aneurysms would likely be an unpalatable option for most male patients. Therefore, we deployed transcriptomic profiling in order to clarify mechanisms downstream of 17 β-estradiol that could be leveraged for future therapies. These global gene expression changes suggest that 17 β-estradiol modulates TNFα mediated NF-κB gene induction of pro-inflammatory proteins that promote a phenotypic switch in aortic SMCs from a contractile to a more inflammatory-like state. This identity switch allows SMCs to promote aortic wall inflammation thus leading to the progression of aneurysm formation. Our in vitro experiments are consistent with this hypothesis as 17 β-estradiol repressed the induction of similar pro-inflammatory genes by TNFα in an NF-κB dependent manner. The exact mechanism of 17 β-estradiol inhibition of NF-κB has been shown to involve several potential avenues including the stability of IκBα, blocking NF-κB nuclear translocation, and the direct inhibition of gene expression on promoters^30-32^. There are many examples of this type of SMC phenotypic switch in vascular disease mediated by TNFα - NF-κB signaling. For example, atherosclerosis can induce the expression of Vcam-1, Icam-1, Cxcl12, and Cx3cl1 in SMCs which in turn facilitates binding to monocytes^33^. TNFα can also induce cerebral SMC phenotypic changes through KLF4 and myocardin which may contribute to cerebral aneurysm formation^34^. In addition, many studies have demonstrated the enhancement of TNFα -NF-κB signaling in AAAs and the presence of modulated SMCs that express inflammatory markers^35^.

In fact, NF-κB inhibition can slow the growth of aneurysms in animal models of AAA^36^. As a result, TNFα -NF-κB signaling might be a universal pathogenic response to injured vascular SMCs, and we are the first to demonstrate the ability of 17 β-estradiol to mitigate this response in Marfan aneurysms.

## Conclusion

In summary we describe the effects of 17 β-estradiol on the reduction of aortic root aneurysms and the incidence of aortic rupture in a Marfan mouse model. Mechanistically, 17 β-estradiol appears to quell inflammation by preventing a TNFα -NF-κB mediated phenotypic switch in SMCs from a contractile to a more inflammatory-like state. These results provide several novel potential targets that could be used therapeutically to treat aortic aneurysms and dissection.

## Supporting information

Cover art

## Abbreviations

TAA: Thoracic aortic aneurysms
AAA: Abdominal aortic aneurysms
SMC: smooth muscle cell
IQR: interquartile range

