## Supplementary figures and images for "17 β-estradiol impedes aortic root dilation and rupture in male Marfan mice"

### Cover art

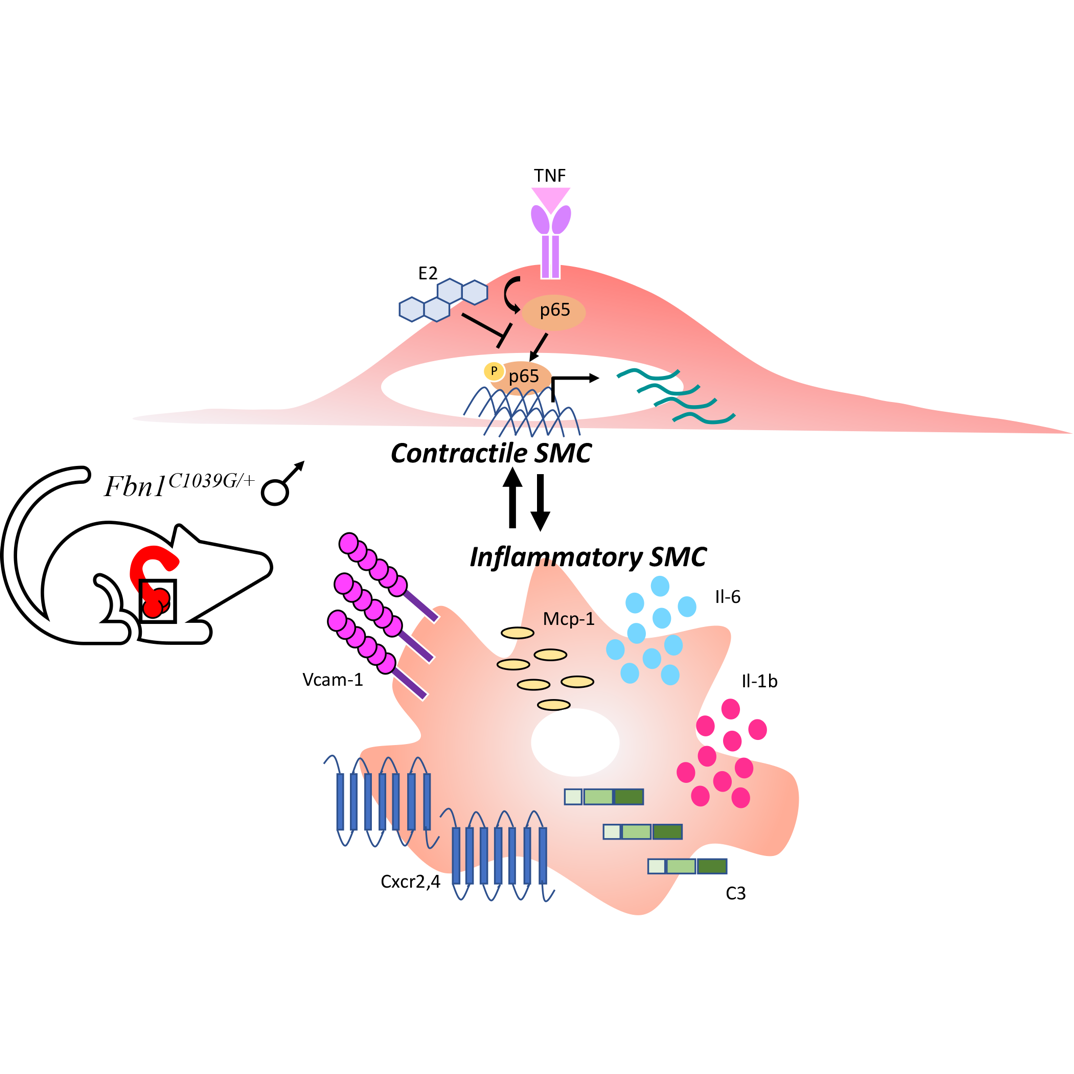

### Supplemental Figures

Supplemental Figure 1

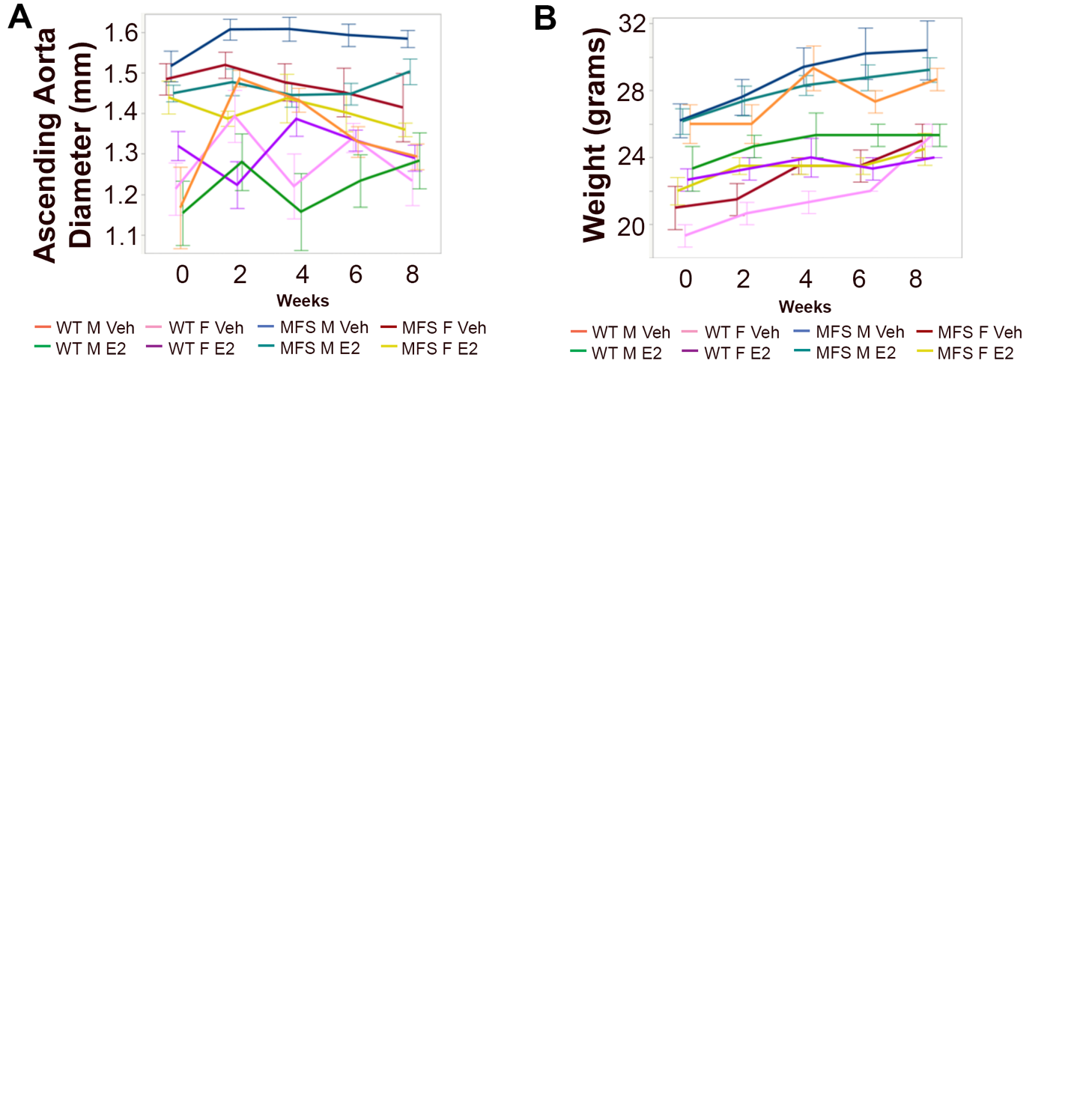

Supplemental Figure 2

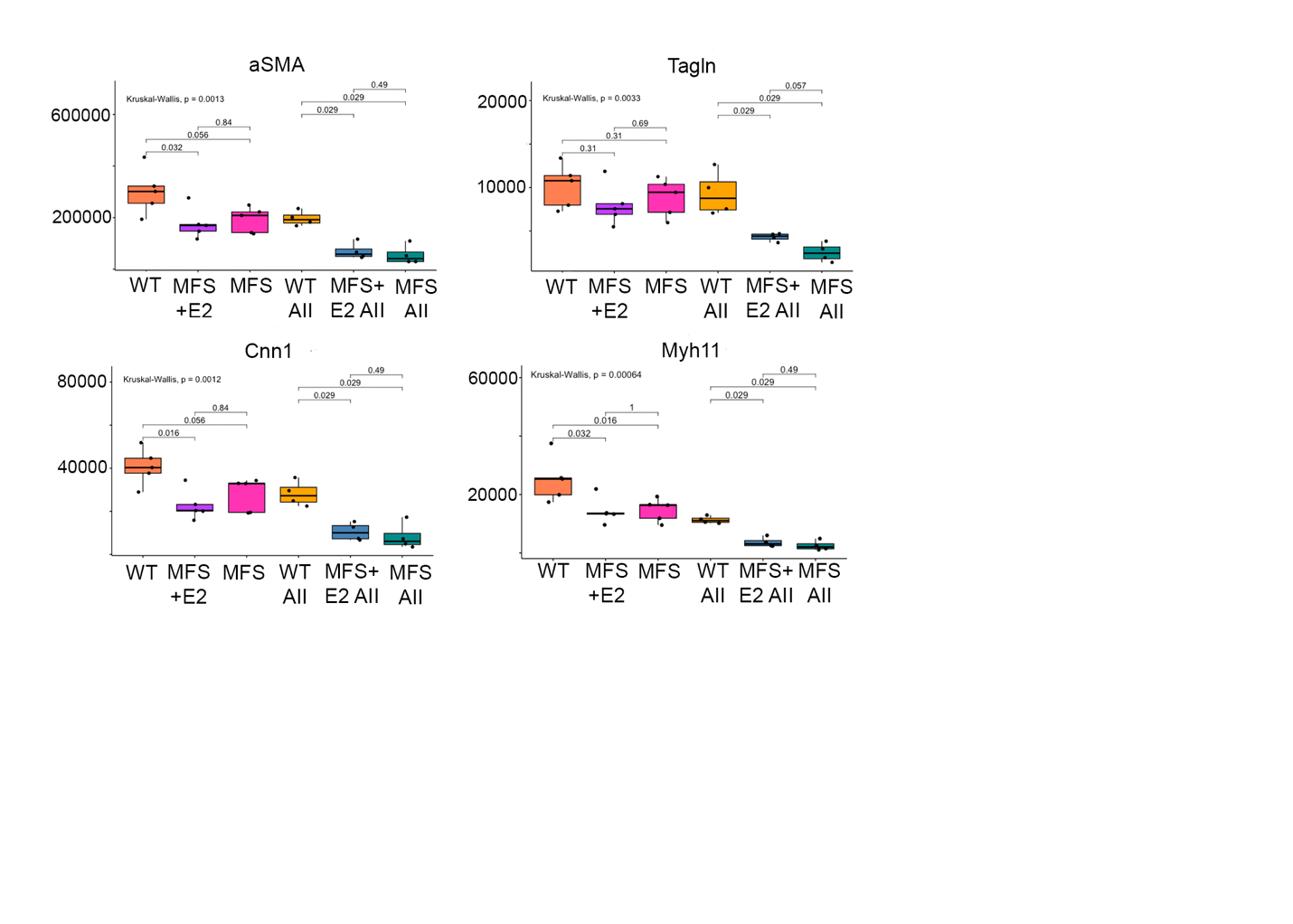
